# Memory reconsolidation impairments in sign-tracking to an audiovisual compound stimulus

**DOI:** 10.1101/2020.03.04.964395

**Authors:** Mohamed L. Drame, Maria Balaet, Jonathan L. C. Lee

## Abstract

Studies of memory reconsolidation of pavlovian memories have typically employed unimodal conditioned stimuli, despite the use of multimodal compound stimuli in other settings. Here we studied sign-tracking behaviour to a compound audiovisual stimulus. First, we observed not unexpectedly that sign-tracking was poorer to the audiovisual compound than to unimodal visual stimuli. Then, we showed that, depending on the parameters of compound stimulus re-exposure at memory reactivation, systemic MK-801 treatment either impaired extinction to improve sign-tracking at test, or disrupted reconsolidation to impair test behaviour. When memory reactivation consisted of re-exposure to only the auditory component of the compound stimulus, we observed sign-tracking impairments following MK-801 treatment, but only under certain test conditions. This was in contrast to the consistent impairment following reactivation with the full audiovisual compound. Moreover, the parameters of auditory stimulus presentation to enable MK-801-induced impairment at test varied depending on whether the stimulus was presented within or outside the training context. These findings suggest that behaviour under the control of appetitive pavlovian compound stimuli can be modulated by targeting both extinction and reconsolidation, and that it is not necessary to re-expose to the full compound stimulus in order to achieve a degree of modulation of behaviour.

## 1. Introduction

Memory reconsolidation is the process hypothesized to take place in order to restabilize a memory that has been destabilized upon reactivation [1]. Pharmacological impairments of memory reconsolidation present as behavioural performance deficits following drug treatment combined with a memory reactivation session [2]. Such experimentally-induced amnesia has been proposed to be a therapeutic strategy for conditions including drug addiction [3, 4].

Reward-associated stimuli acquire motivational incentive value through appetitive learning processes, which subsequently contribute to relapse-like behaviour [5]. Stimuli with incentive value elicit approach that can be evaluated as sign-tracking behaviour [6]. Sign-tracking paradigms in rodents traditionally use simple visual stimuli alone [7], and reconsolidation deficits have been observed in such settings [8, 9]. However, while more translationally-relevant compound stimuli have been studied in fear memory reconsolidation studies, showing that re-exposure to one component triggers reconsolidation of the other element [10], it remains unclear whether such principles of memory destabilization also apply in appetitive settings.

Here we explored sign-tracking to compound audiovisual stimuli associated with sucrose reward. As tests of sign-tracking require an index of truly discriminated approach between concurrently-presented stimuli [7], this necessitated presentation of the visual element of the compound stimulus alone, but we also tested approach behaviour in response to the intact compound in a serial stimulus presentation test [9]. In such a paradigm, we first established the parameters of compound stimulus re-exposure necessary to destabilize the incentive CS-sucrose memory. We used the NMDA receptor antagonist MK-801 as the amnestic agent as we and others have shown that systemic injections of MK-801 impair reconsolidation across a number of settings [11–18], including sign-tracking [8, 19]. We then were able to test the hypothesis that elemental stimulus presentation can destabilize a compound pavlovian appetitive stimulus. Moreover, we tested the translationally-relevant question of whether elemental stimulus presentation outside the original context of conditioning (akin to in a clinical treatment setting) is able to trigger incentive memory destabilization.

## 2. Methods

### 2.1. Animals

232 male Lister Hooded rats (Charles River, UK; 200-250 g at the start of the experiment) were housed in quads under a 12 h light/dark cycle (lights on at 0700) at 2I°C with water provided ad libitum apart from during the behavioural sessions. Standard cages contained aspen chip bedding and environmental enrichment was available in the form of a Plexiglass tunnel. The rats were food restricted and fed 80 g/cage/day chow from the first day of behavioural training. Experiments took place in a behavioural laboratory between 0830 and 1300. At the end of the experiment, animals were humanely killed via a rising concentration of CO2; death was confirmed by cervical dislocation. Principles of laboratory animal care were followed, as approved by the University of Birmingham Animal Welfare and Ethical Review Body and in accordance to the United Kingdom Animals (Scientific Procedures) Act 1986, Amendment Regulations 2012 (PPL P8B15DC34 and P3B19D9B2).

### 2.2. Drugs

MK-801 was administered intra-peritoneally at a previously-established dose (0.1 ml/kg; 0.1 mg/ml in saline) immediately after the reactivation session [14]. Allocation to drug treatment was fully randomised within each experimental cohort of 16 or 32 rats.

### 2.3. Behavioural procedures

Behavioural training and testing took place in operant chambers (Med-associates, VT) as previously described [20]. On the first day of behavioural training, rats were pretrained to be familiarised to the sucrose pellet reward. 45-mg sucrose pellets were delivered on a random schedule (interval of 4 – 79 s) in a 20-min session. Starting the next day and daily for 10 consecutive weekdays, rats were trained to associate a 10-s audiovisual compound stimulus with delivery of a single sucrose pellet (at the end of the stimulus presentation). The compound stimulus consisted of (1) extension of the left lever, illumination of the left stimulus light and presentation of a 3-kHz tone, or (2) extension of the right lever, illumination of the right stimulus light and presentation of a 1-Hz clicker. The reinforcement of stimulus (left/tone vs right/clicker) was randomised across rats. There was a random interval (9 – 39 s) between stimulus presentations, and the order of stimulus presentations was randomised within each consecutive pair of presentations. Each session consisted of 50 stimulus presentations (25 of each stimulus).

Memory reactivation varied across behavioural conditions, and took place 2 d after the end of training. However, stimulus presentation (either the full audiovisual compound or just the auditory stimulus) followed the same schedule as during training. Sucrose reward was never delivered during reactivation. For experiments, in which the auditory stimulus was presented outside the training context, rats were brought into the testing room and placed individually in fresh standard cages. A single operant chamber was used, with the sound-attenuating chamber left open, and the auditory stimuli presented as per the preceding experiments.

On the day after memory reactivation, the first test was conducted. For sign-tracking rats, the first test consisted of a probe test, in which the two visual stimuli were simultaneously presented (no auditory stimuli were presented), again on the same schedule as for training (random 9-39-s interval), with a total of 20 concurrent stimulus presentations. For the goal-tracking rats, the test consisted of the same experience as a training session, but with no sucrose pellet delivery. Sign-tracking rats received this test 3 days after the probe test. Goal-tracking rats just had the single test.

### 2.4. Statistical analysis

Rats were first allocated as sign-trackers or goal-trackers. Sign-tracking rats displayed discriminated approach to the CS+ on the final day of training; determined by automatic contact with the extended lever upon approach [8, 19, 21], This was quantified as a numerically greater number approaches to the CS+ compared to the CS-, and approach to the CS+ on at least 50% of CS+ presentations. Goal-tracking rats were those rats that were not classified as sign-trackers, and which displayed a greater number of nosepoke entries to the sucrose magazine during the CS+ compared to the CS-. Rats that showed neither discriminated CS+ approach not discriminated CS+ nosepoking were not included in either group and were not tested or analysed further (n=12 across all experiments).

Data are presented as mean number (+ SEM) of approaches to the CS+/CS- or mean number of nosepokes during the CS+/CS-at test and memory reactivation. Data were analysed in JASP 0.9.1.0 [22] using repeated measures ANOVA with CS and MK-801 as factors, with alpha=0.05 and η^2^_p_ reported as an index of effect size, and the data were initially checked for normality. The data were also checked for sphericity, and the Greenhouse-Geisser correction applied as appropriate. Significant effects of MK-801 or MK-801 x CS interactions were followed by analyses of simple main effects of MK-801 on approach to/nosepoking during each CS.

## 3. Results

### 3.1. Sign-tracking vs goal-tracking

First, we established sign-tracking with a compound audiovisual stimulus. The visual stimulus was the insertion of a lever and the illumination of a stimulus light above the lever. The compounded auditory stimulus was either a tone or clicker. Compared to previous studies using identical training parameters using only the visual components of the current stimuli [8], sign-tracking behaviour was observed in a smaller proportion of rats. Approximately 43% of rats showed sign-tracking behaviour, approaching the CS lever as the first response to CS presentation, and showing discriminated lever approach between the CS+ and CS-. The majority of the remaining rats appeared to show goaltracking behaviour, nosepoking first and showing discriminated nosepoking, rather than discriminated lever approach. An analysis of our initial experimental group (which later received 50 CS presentations at reactivation) revealed that those rats designated as sign-trackers (see methods) showed greater discriminated lever approach (Fig. 1A; Group x CS x day: F(5.0,199.5)=31.0, p<0.001, η2_p_=0.44; Group x CS: F(1,40)=143.8, p<0.001, η2_p_=0.78) and approached the CS+ lever first more (Fig. 1C; Group x day: F(3.7,147.5)=31.4, p<0.001, η2_p_=0.44; Group: F(1,40)=163.8, p<0.001, η2_p_=0.80) than non-sign-trackers through training. Furthermore, sign-trackers showed less discriminated nosepoking (Fig. 1B; Group x day: F(4.6,185.0)=10.1, p<0.001, η2_p_=0.20; Group: F(1,40)=21.4, p<0.001, η2_p_=0.35) and nosepoked first less (Fig. 1D; Group x day: F(4.5,180.7)=27.5, p<0.001, η2_p_=0.41; Group: F(1,40)=88.2, p<0.001, η2_p_=0.69) than non-sign-trackers.

**Figure 1.**
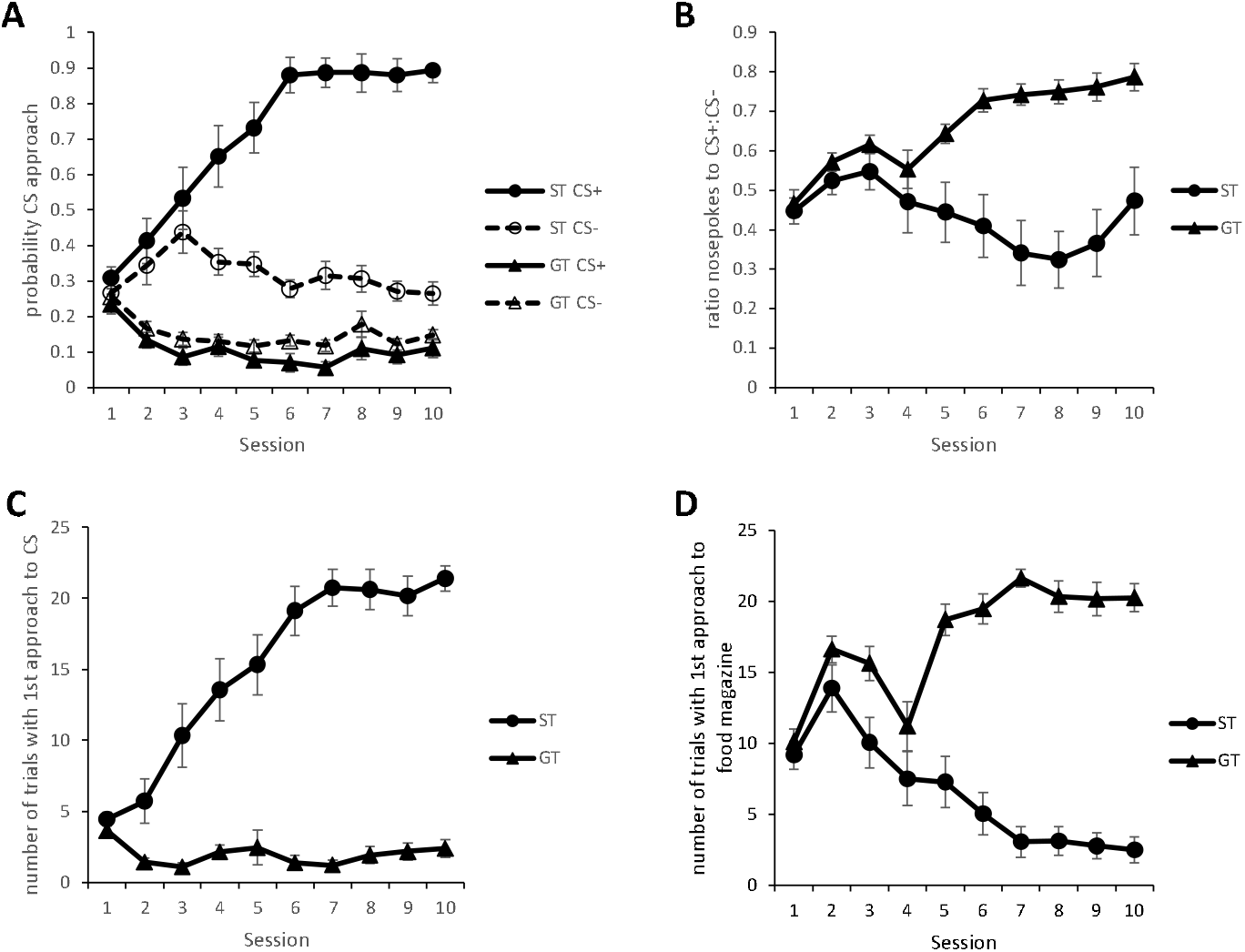
Emergence of sign-tracking and goal-tracking to compound audiovisual stimuli in different rats. Sign-tracking rats showed greater discriminated approach to the CS+than did goal-tracking rats (**A**). Goal-tracking rats showed a greater ratio of nosepokes to the CS+:CS-than did sign-tracking rats (**B**). Upon CS+ presentation, sign-tracking rats tended to approach the CS first (**C**), whereas goal-tracking rats preferentially nosepoked in the food magazine first (**D**). Data presented as mean ± SEM. N=18 sign-trackers & 24 goal-trackers.

### 3.2. Effects of post-reactivation MK-801 on Sign-tracking

Sign-tracking rats were subjected to varying compound CS re-exposure at memory reactivation, receiving an i.p. injection of MK-801 immediately after reactivation. Using our previously-effective reactivation parameters of 50 CS presentations (Lee & Everitt, 2008), there was some evidence for an impairment of extinction, rather than reconsolidation. A main effect of MK-801 was observed at an initial probe test of discriminated approach to the visual stimuli alone (Fig. 2A; F(1,16)=5.73, p=0.029, η^2^_p_=0.26), with no MK-801 x CS interaction (F(1,16)=2.34, p=0.15, η^2^_p_=0.13). Analysis of simple main effects revealed an effect of MK-801 on CS+ approach (F=10.1, p=0.006), but not CS-approach (F=0.045, p=0.84). These differences at test were not attributable to pre-treatment group differences, as performance at the reactivation session was not statistically different between the groups (data not shown; MK-801 x CS: F(1,16)=0.037, p=0.85, η^2^_p_=0.002, BF10=0.44). At a subsequent test of approach to the compound CS (Fig. 2B), there was no effect of MK-801 (F(1,16)=0.60, p=0.45, η^2^_p_=0.036), or MK-801 x CS interaction (F(1,16)=0.011, p=0.92, η^2^_p_=0.001).

**Figure 2.**
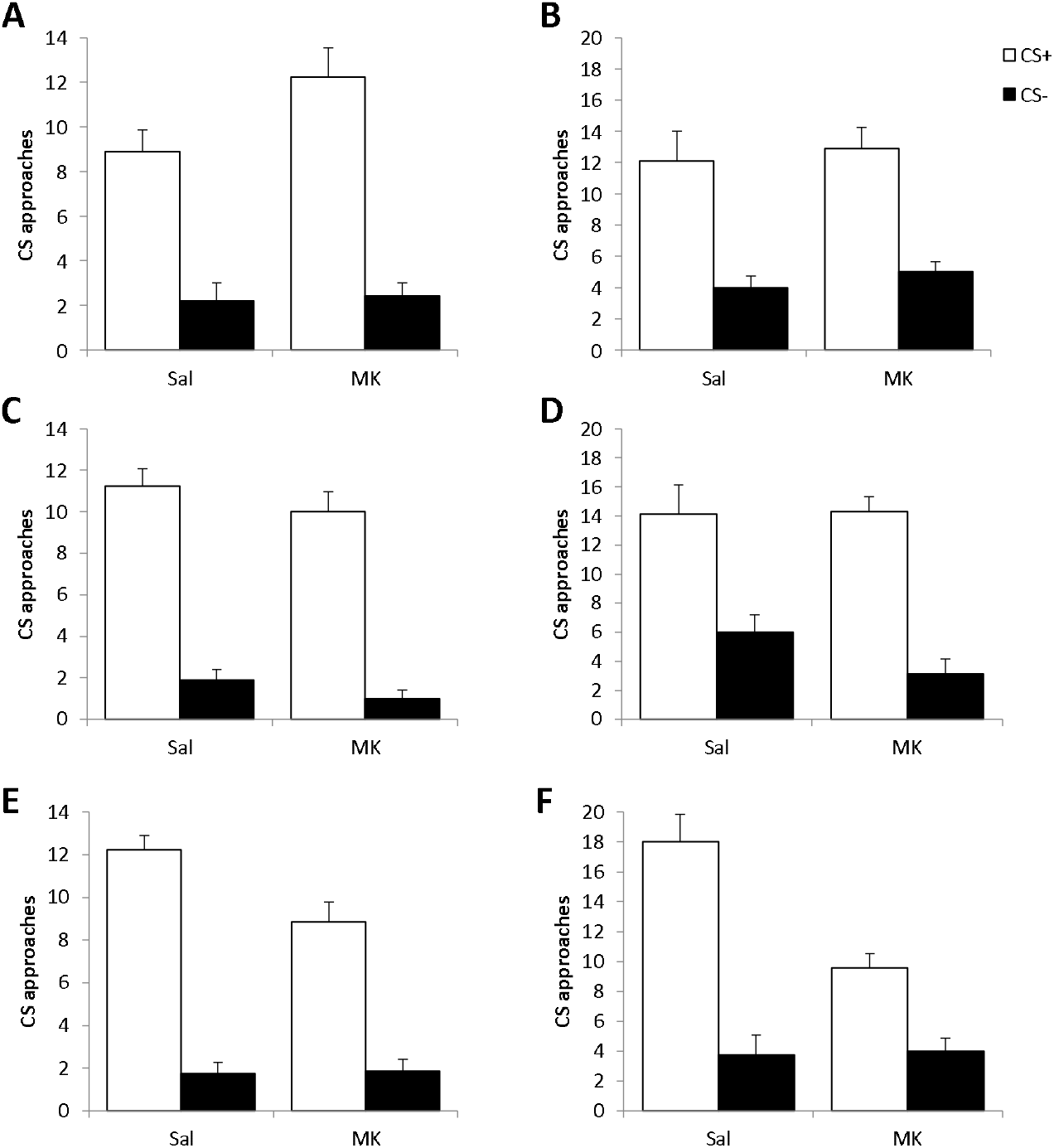
Sign-tracking following memory reactivation with the compound audiovisual stimulus. Following reactivation with 50 CS re-exposures, MK-801-treated rats approached the CS+ more than Saline-treated controls at the initial probe test (**A**), but not at the subsequent test of approach to the compound stimuli (**B**). With 20 CS presentations at reactivation, there was no effect of MK-801 at either the probe test (**C**) or the compound stimuli test (**D**). When memory reactivation consisted of 10 CS presentations, MK-801-treated rats showed impaired approach to the CS+ at both the probe test (**E**) and the test of approach to the compound stimuli (**F**). Data presented as mean + SEM. N=7-9 per group.

Given the complex relationship between reconsolidation and extinction, we reduced the number of CS re-exposures at reactivation to 20. Under these conditions, there was no evidence for an effect of MK-801 on either reconsolidation or extinction. At the initial probe test (Fig. 2C), there was no main effect of MK-801 (F(1,14)=2.89, p=0.11, η^2^_p_=0.17) or MK-801 x CS interaction (F(1,14)=0.051, p=0.83, η^2^_p_=0.004). Similarly, there was no main effect of MK-801 (F(1,14)=0.66, p=0.43, η^2^_p_=0.045) or MK-801 x CS interaction (F(1,14)=2.01, p=0.18, η^2^_p_=0.13) at the subsequent test of approach to the compound CS (Fig. 2D).

As there is evidence for intermediate parameters that are ineffective at engaging NMDAR-dependent reconsolidation or extinction [23, 24], we further reduced CS re-exposure to 10 presentations at reactivation. Under these conditions, there was evidence for an MK-801-induced impairment in discriminated approach to the CS. At the probe test (Fig. 2E), there was an MK-801 x CS interaction (F(1,13)=6.19, p=0.027, η^2^_p_=0.32) as well as a main effect of MK-801 (F(1,13)=11.53, p=0.005, η^2^_p_=0.47). Analysis of simple main effects revealed an effect of MK-801 on CS+ approach (F=24.6, p<0.001), but not CS-approach (F=0.025, p=0.88). At the subsequent test of approach to the compound CS (Fig. 2F), there remained an MK-801 x CS interaction (F(1,13)=12.30, p=0.004, η^2^_p_=0.49) as well as a main effect of MK-801 (F(1,13)=13.24, p=0.003, η^2^_p_=0.51). Analysis of simple main effects again confirmed an effect of MK-801 on CS+ approach (F=28.1, p<0.001), but not CS-approach (F=0.025, p=0.88). This was also reflected in a reduction in the number of test trials in which the rats approached the CS+ first during its presentation (Saline=17.0±2.2, MK-801=9.3±1.0; F(1,13)=13.1, p=0.003, η^2^_p_ =0.50). These differences at test were not attributable to pre-treatment group differences, as performance at the reactivation session was not statistically different between the groups (data not shown; MK-801 x CS: F(1,13)=0.17, p=0.68, η^2^_p_=0.013; MK-801: F(1,13)=3.48, p=0.085, η^2^_p_=0.21).

An exploratory analysis of all reactivation conditions confirmed that the effect of MK-801 depended upon the number of CS re-exposures at reactivation (Re-exposures x MK-801 x CS: F(2,43)=3.76, p=0.031, η2_p_=0.15).

Next we explored whether exposure to the auditory element of the compound stimulus alone was able to destabilise the sign-tracking memory. 10 exposures to the auditory stimulus did not replicate the effect of 10 exposures to the full compound stimulus, with MK-801 having no effect on subsequent sign-tracking behaviour to the visual stimulus at the probe test (Fig. 3A; MK-801: F(1,17)=0.10, p=0.75, η^2^_p_=0.006; MK-801 x CS: F(1,17)=0.003, p=0.96, η^2^_p_=0.000). Similarly, there was no main effect of MK-801 (F(1,17)=0.02, p=0.90, η^2^_p_=0.001, BF10=0.40) or MK-801 x CS interaction (F(1,17)=0.06, p=0.82, η^2^_p_=0.003, BF10=0.43) at the subsequent test of approach to the compound CS (Fig. 3B).

**Figure 3.**
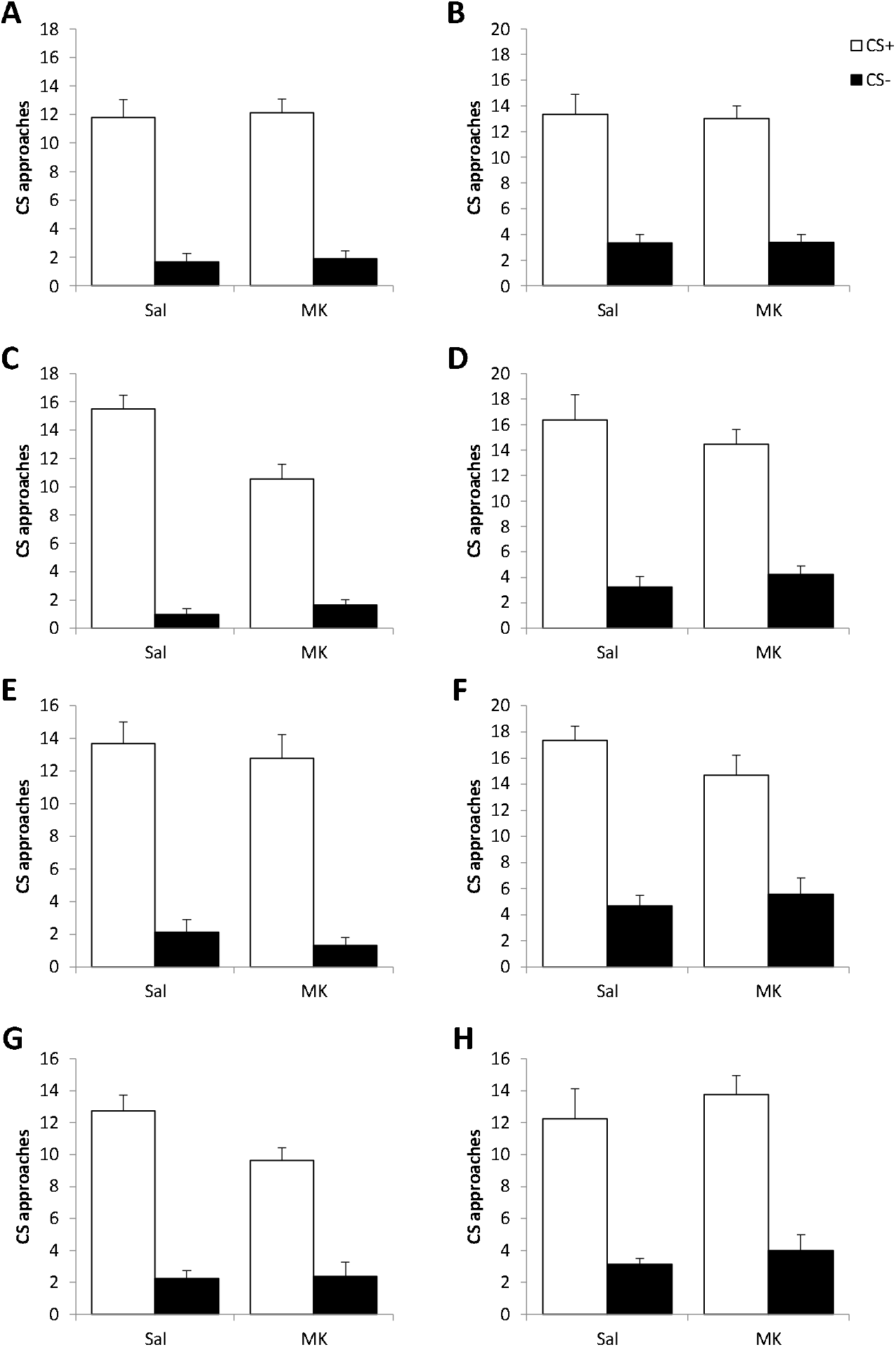
Sign-tracking following memory reactivation with the auditory element of the compound stimulus. Following reactivation with 10 auditory CS re-exposures in the training context, there was no effect of MK-801 at the initial probe test (**A**) or at the subsequent test of approach to the compound stimuli (**B**). With 20 auditory CS presentations at reactivation in the training context, MK-801-treated rats showing impaired CS+ approach at the probe test (**C**), but not at the compound stimuli test (**D**). When rats experienced the auditory stimulus presentation outside the training context at memory reactivation, MK-801 treatment following 20 presentations resulted in no effect at the probe test (**E**) or at the test of approach to the compound stimuli (**F**). When the auditory stimuli were presented 50 times outside the training context, MK-801 treatment did result in impaired CS+approach at the probe test (**G**), but not at the compound stimuli test (**H**). Data presented as mean + SEM. N=7-10 per group.

As it is possible that presentation of the auditory stimulus alone is less salient and might be insufficient to trigger memory destabilisation, we increased the number of presentations to 20. Under these conditions MK-801 did impair subsequent sign-tracking behaviour to the visual stimulus at the probe test (Fig. 3C; MK-801: F(1,15)=8.21, p=0.012, η^2^_p_=0.35; MK-801 x CS: F(1,15)=16.3, p=0.001, η^2^_p_=0.52). Analysis of simple main effects confirmed an effect of MK-801 on approach to the CS+ (F=21.9, p<0.001), but not on CS-approach (F=0.40, p=0.54). However, this disruptive effect of MK-801 did not persist to the test of approach to the compound CS (Fig. 3D; MK-801: F(1,15)=0.18, p=0.68, η^2^_p_=0.012; MK-801 x CS: F(1,15)=1.59, p=0.23, η^2^_p_=0.096). There was also little evidence for a difference in the number of test trials with rats approaching the CS+ first (Saline=15.8±1.8, MK-801=13.3±1.3; F(1,15)=1.38, p=0.26, η^2^_p_=0.084). In order to determine whether the lack of effect at the second test was due to the prior test experience or the delay from reactivation to test, we repeated the experiment proceeding directly to the test of approach to the compound CS. There remained no effect of MK-801 under these test conditions (data not shown; MK-801: F(1,16)=0.76, p=0.40, η^2^_p_=0.046; MK-801 x CS: F(1,16)=1.40, p=0.25, η^2^_p_=0.080).

Finally, we evaluated the capacity of presentation of the auditory stimulus outside the training context to trigger memory destabilization. Rats were brought to the testing room, but remained outside the testing chambers while the auditory stimuli were presented. With 20 stimulus presentations, MK-801 did not affect subsequent performance at the probe test (Fig. 3E; MK-801: F(1,16)=1.47, p=0.24, η^2^_p_=0.084; MK-801 x CS: F(1,16)=0.002, p=0.97, η^2^_p_=0.000) or at the test of approach to the compound CS (Fig. 3F; MK-801: F(1,16)=0.53, p=0.48, η^2^_p_=0.032; MK-801 x CS: F(1,16)=2.80, p=0.11, η^2^_p_=0.15). As the presentation of auditory stimuli outside the testing chamber may also be less salient, we increased the number of stimulus presentations to 50. Under these conditions, MK-801 did produce an impairment sign-tracking behaviour to the visual stimulus at the probe test (Fig. 3G; MK-801: F(1,15)=5.65, p=0.031, η^2^_p_=0.27; MK-801 x CS: F(1,15)=4.96, p=0.042, η^2^_p_=0.25). Analysis of simple main effects confirmed an effect of MK-801 on approach to the CS+ (F=8.88, p=0.009), but not on CS-approach (F=0.022, p=0.89). Once again, the effect of MK-801 did not persist to the test of approach to the compound CS (Fig. 3H; MK-801: F(1,15)=0.36, p=0.56, η^2^_p_=0.023; MK-801 x CS: F(1,15)=0.042, p=0.84, η^2^_p_=0.003; effect of MK-801 on trials with approach to the CS+ first: Saline=11.0±2.1, MK-801=12.9±1.2; F(1,15)=0.19, p=0.67, η^2^_p_=0.012).

### 3.3. Effects of post-reactivation MK-801 on Goal-tracking

Goal-tracking rats were also subjected to a variety of reactivation conditions. Following reactivation consisting of 20 presentations of the compound stimulus, there was evidence for an MK-801-induced increase nosepokes to the CS+ following 20 stimulus presentations (Fig. 4A; MK-801: F(1,16)=1.35, p=0.26, η^2^_p_=0.078; MK-801 x CS: F(1,16)=5.80, p=0.028, η^2^_p_=0.27), indicating an impairment of extinction. However, there was no effect on the number of CS+ trials on which rats first performed a nosepoke (Saline=6.6±1.1, MK-801=7.0±1.5; F(1,16)=0.36, p=0.56, η^2^_p_=0.022). Following reactivation with 10 presentations of the compound stimulus, there was little evidence for any effect of MK-801 (Fig. 4B; MK-801: F(1,15)=0.47, p=0.50, η^2^_p_=0.030; MK-801 x CS: F(1,15)=1.62, p=0.22, η^2^_p_=0.098; effect of MK-801 on trials with a nosepoke first: Saline=6.6±1.3, MK-801=10.1±1.9; F(1,15)=2.82, p=0.11, η^2^_p_=0.16). With a further reduction in stimulus presentation at reactivation to 4, MK-801 still had no effect on nosepoking (Fig. 4C; MK-801: F(1,22)=0.096, p=0.76, η^2^_p_=0.004; MK-801 x CS: F(1,22)=1.44, p=0.24, η^2^_p_=0.062), with weak evidence for a reduction in the number of trials at which the rats nosepoked first (Saline=8.6±1.5, MK-801=5.0±1.3; F(1,22)=3.69, p=0.068, η^2^_p_=0.14). Therefore, there was little evidence for an impairment of goal-tracking in the current experimental setting.

**Figure 4.**
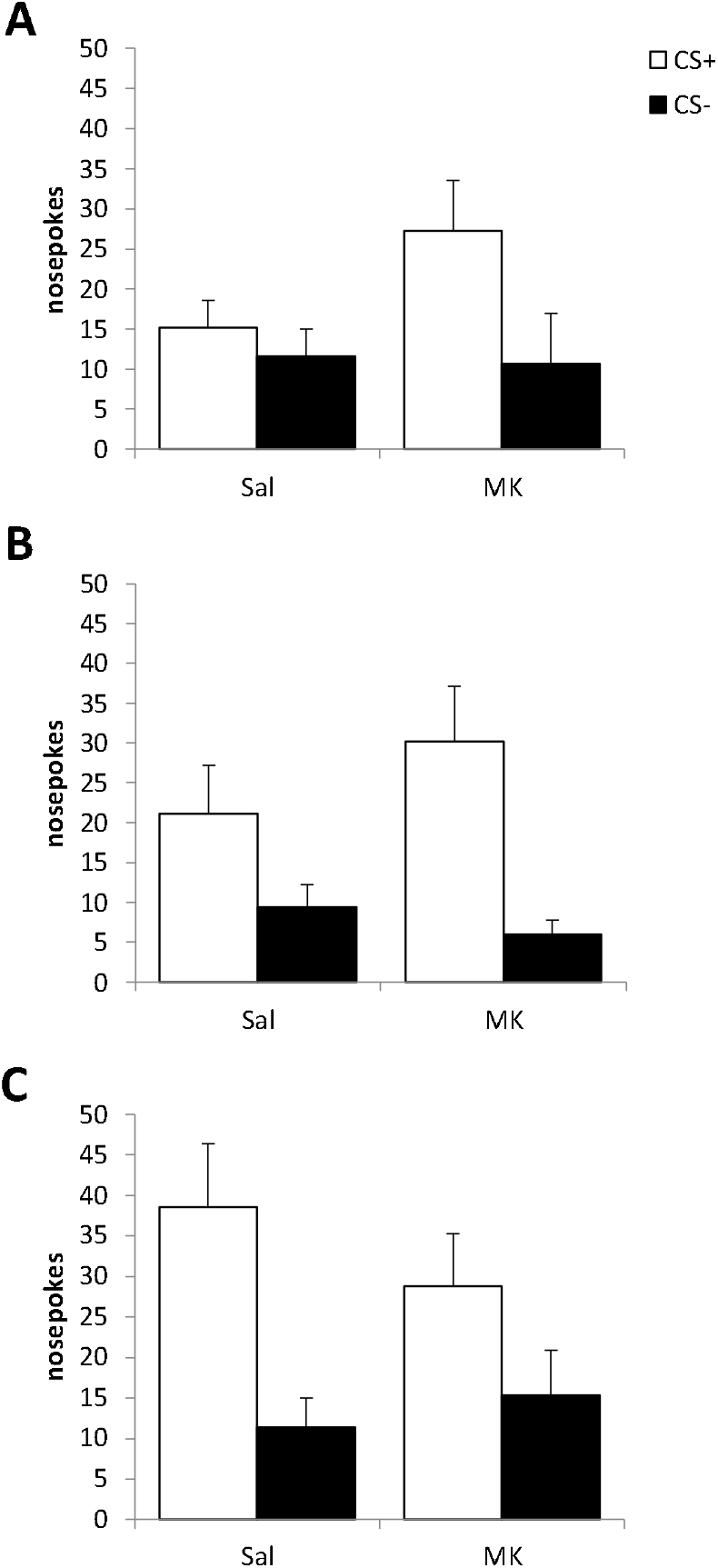
Goal-tracking to a compound audiovisual stimulus. Following reactivation with 20 CS presentations, MK-801-treated rats nosepoked more than Saline-treated controls during the goal-tracking test (**A**). With 10 CS presentations at memory reactivation, there was no effect of MK-801 at the subsequent test (**B**). When memory reactivation consisted of 4 CS presentations, there was little evidence for an impairment of goal-tracking at test (**C**). Data presented as mean + SEM. N=8-12 per group.

## 4. Discussion

Our current data show that the appetitive pavlovian association underpinning sign-tracking to a compound audiovisual stimulus destabilises under selective parametric conditions at memory reactivation. We also demonstrate that presentation of only the auditory component of the compound stimulus at memory reactivation can trigger memory destabilisation, both when presentation takes place inside and outside the training context. However, the parameters of effective stimulus presentation differ depending on the method of presentation. Finally, rats that did not acquire sign-tracking during training typically displayed goal-tracking behaviour, for which there was no evidence for impaired reconsolidation in the present behavioural paradigm.

While sign-tracking is typically conducted using a simple or compound visual stimulus [7, 21], especially for studies of memory reconsolidation [8, 9], compound audiovisual stimuli have also been employed [e.g. 25]. In our previous study of memory reconsolidation in a sign-tracking setting [8], 33 out of 48 rats successfully acquired sign-tracking behaviour using the same criteria as employed here. In the present study, the ratio of sign-trackers to goal-trackers was closer to 50:50 across all experiments. This is likely due to the fact that sign-tracking is more likely to be acquired to highly localizable visual stimuli [26, 27], whereas goal-tracking is promoted by auditory stimuli [26]. Therefore, there may be individual differences in the salience attributed to, and the attention directed towards, the visual and auditory elements of our compound stimuli. It is possible that those rats that attend more to the visual component will develop sign-tracking, whereas rats whose attention is directed more to the auditory component show goal-tracking behaviour. Such individual differences may be related to the emerging differential mechanisms of sign-tracking and goal-tracking behaviour [e.g. dopamine D2R activity; 26].

In our sign-tracking rats, we demonstrated a pattern of results similar to those previously observed in appetitive and aversive memories [13, 23, 24], in that while there are parameters of stimulus reexposure that promote memory destabilisation and extinction, there is an intermediate level of reexposure, at which MK-801 treatment has no observable effect. As also appears to be the case for contextual fear memories [13], it is difficult to determine *a priori* the effective parameters for triggering memory destabilisation or extinction. While 50 stimulus re-exposures engaged extinction in the present study, the same number resulted in destabilisation previously [8]. It remains unclear whether this difference is due to the compound vs simple stimuli used in the two studies, or to some other as yet unidentified factor. For example, in our previous study [8], the reinforcer was liquid sucrose (as opposed to sucrose pellets here).

Notwithstanding the overall similarity to previous studies above, the use of two different tests in the current study presents additional insight into the behavioural effects of MK-801. While there was an increase in discriminated approach to the CS+ in the visual probe test under the 50-CS reactivation condition, there was no difference between MK-801 and Saline rats at the subsequent compound stimulus test. There are three potential interpretations of this apparent recovery. First, it may simply reflect the expected spontaneous recovery that follows normal extinction training [28], such that the saline-administered control rats recovered responding at the second test, conducted 3 days after the first test. With heroin reward, extinction of sign-tracking readily recovered at a second test, albeit after a more substantial delay of 3 weeks [29]. There is no reason why the MK-801-treated rats that showed impaired extinction at the probe test should elevate their responding at the second test, and so spontaneous recovery would bring performance in the two groups to equivalent levels. However, the tests differ in terms of the presence of the auditory component of the compound stimulus. Therefore, it is possible that the addition of the auditory component recovered responding at the second test. However, given that it was the full compound stimulus that was presented during extinction training, it is unlikely that performance deficit would be observed only when the test involved presentation of the visual component. Finally, the simultaneous vs sequential nature of CS+/CS-presentation in the two tests may be an important factor. The probe test truly evaluates discriminated approach to the CS+ in a choice test [7, 21], and so it is possible that extinction of the CS+ is only observable under probe test conditions. Indeed, when test does not involve discrimination between a CS+ and CS-, extinction of sign-tracking following equivalent training parameters is generally not observed [25, 30].

When the reactivation session involved re-exposure to only the auditory component of the compound stimulus, the number of stimulus presentations necessary to induce successful memory destabilization was greater than required with compound stimulus presentation. While we are not aware of equivalent observations in different (e.g. fear memory) settings, extinction of sign-tracking in pigeons is more effective when it involves non-reinforced compound stimulus presentation, as compared to presentation of a single element [31; but note that the elements of the compound stimulus were conditioned separately]. There was an even greater stimulus re-exposure requirement when the rats were re-exposed outside the training context, as compared to inside the operant chambers. This may be due to the contextual discrepancy between conditioning and reactivation. For example, the change in context of presentation may influence the degree of prediction error engaged by a particular number of stimulus presentations, such that the threshold of prediction error required to stimulate memory destabilization [32, 33] is only reached by a higher number. Alternatively, there may also be competition between new learning and reconsolidation-updating. We have shown previously that consolidation of new learning and reconsolidation-mediated updating of existing memories engage doubly dissociable mechanisms in a contextual fear memory setting [34], It may be that a greater number of elemental stimulus presentations is required to induce memory destabilization outside the training context, in order to overcome the tendency to encode new memories in a novel context [35]. If this were the case, we would predict that there would similarly be a requirement for more compound stimulus presentations outside the training context, compared to the 10 required inside the operant chamber. However, regardless of the explanation for the parametric differences, the observation that partial stimulus re-exposure in a setting closer to the home cage than the conditioning chambers (albeit in the same experimental room) is sufficient to destabilize the incentive memory underpinning sign-tracking has translational relevance for the application of reconsolidation-based treatment strategies for problematic reward-seeking behaviours [e.g. 36].

For the apparent reconsolidation deficit in sign-tracking, there was a performance deficit in MK-801-treated rats across both tests. This shows the persistence of the deficit, as is important for the demonstration of reconsolidation impairment [37] and that it generalises across test settings. While such a persistent deficit was observed following reactivation with the compound stimulus, it was not evident when reactivation involved exposure to the auditory component alone, whether or not such presentation occurred inside or outside the training context. First, it is important to note that reactivation of the appetitive memory through re-exposure to just the auditory element of the compound stimulus must have destabilised the entire compound CS-sucrose association, as discriminated approach to the visual component was impaired. This is consistent with findings from a study on fear memory reconsolidation, in which intra-amygdala anisomycin following reactivation through re-exposure to the visual component resulted in subsequent impaired conditioned freezing to the auditory component [10]. However, the study by Debiec et al (2013) did not test conditioned freezing to the full compound stimulus. Here, the apparently normal responding to the compound stimulus likely again reflects the difference in the test procedures, rather than any recovery of performance, especially as omitting the probe test resulted in similarly unimpaired performance in the compound stimulus test when reactivation consisted of the 20 auditory stimulus presentations in the training context. This also rules out an alternative interpretation based upon MK-801-impaired unovershadowing of the visual CS component following “brief” extinction of the auditory component at reactivation. The most likely account for the difference in effects at the two tests is the requirement for CS+/CS- discrimination at the probe test. There is evidence in the literature that such a difference determines whether or not apparent reconsolidation impairments are observed. With propranolol treatment, there was no evidence for an impairment in a discrimination probe test [8], but there was in a setting that involved no control CS- [9]. Therefore, probe discrimination and simple CS approach are not equivalent assays of sign-tracking behaviour. In the absence of a strong alternative, perhaps the most parsimonious interpretation of our results is that presentation of the auditory component did not destabilise the memory as fully as does presentation of the full compound stimulus, leading to impaired performance only under certain test settings.

For our goal-tracking results, we observed no evidence of a disruptive effect of MK-801 under any parameters of compound stimulus re-exposure. This was in spite of a clear effect of MK-801 to elevate goal-tracking following 20 CS presentations at reactivation, which highly likely reflects an impairment of extinction [38]. While it is possible that we simply failed to find the appropriate parameters of CS re-exposure to destabilise the goal-tracking memory trace, this is somewhat unlikely given the range of parameters tested. Nevertheless, reduction to a single presentation each of the CS+ and CS-may have resulted in successful memory destabilization. Alternatively, it is possible that impairments in goal-tracking reconsolidation are less likely to be observed under the current experimental set-up compared to our previous-used paradigm [38]. Previously, we demonstrated an impairment in goal-tracking memory reconsolidation when conditioning occurred to unimodal auditory stimuli [38, 39] as is commonly employed to promote goal-tracking behaviour [25]. Moreover, our measure of goal-tracking in the current study was open to confound as it did not correct for the baseline rates of nosepoking in the interstimulus intervals, which have the potential randomly to vary across the test session and differentially so prior to CS+ and CS- presentation. Therefore, the failure to demonstrate a reconsolidation deficit in goal-tracking in the current study cannot be interpreted as a fundamental inability to disrupt the underpinning memory.

In summary, appetitive sign-tracking memory reconsolidation impairments can be observed with audiovisual compound conditioned stimuli. Memory destabilisation can be induced by exposure to the auditory component of the compound stimulus, although the nature of the reconsolidation deficit may be quantitatively or qualitatively different. Therefore, it is possible to diminish the impact of appetitive compound stimuli on behaviour via reconsolidation impairments without the need to re-expose to the full compound stimulus. These findings support the potential application of reconsolidation-based therapeutic strategies to maladaptive reward-seeking behaviour.

## 5. Acknowledgements

The authors would like to thank David Barber for technical support. The work was supported by grants from the UK BBSRC (BB/J014982/1) and MRC (MR/M017753/1).

